# Handedness and its association with education and spatial navigation assessed in over 400,000 participants across 41 countries

**DOI:** 10.1101/2023.03.30.534904

**Authors:** P. Fernandez-Velasco, A. Coutrot, H. Oloye, J.M. Wiener, R.C. Dalton, C. Hoelscher, E. Manley, M. Hornberger, H.J Spiers

## Abstract

There is an active debate concerning the association of handedness and spatial ability. Past studies used small sample sizes within a single country. Determining the effect of handedness on spatial ability requires a large, cross-cultural sample of participants, and a navigation task with real-world validity. Here, we overcome these challenges via the mobile app Sea Hero Quest. We analysed the navigation performance from 422,772 participants from 41 countries and found no reliable evidence for any difference in spatial ability between left- and right-handers across all countries. Using 749,037 participants from the larger sample, we replicate previous findings that age, gender, and country of residence have an impact on the prevalence of left-handedness, and found an effect of education on left-handedness prevalence in China, Indonesia, India, Taiwan, and Hong Kong. Our study clarifies the factors associated with spatial ability and outlines new ways in which cultural patterns influence handedness.

**Statement of Relevance:** What is the relation between handedness and navigation ability? Evidence so far has been mixed, and findings from small-scale and large-scale tasks seem to point in opposite directions. Part of the reason is that cultural and sociodemographic differences have a significant impact on both spatial ability and handedness. Tackling the question requires a large, cross-cultural sample of participants performing an ecologically valid navigation task. Here, we employ a mobile app, Sea Hero Quest, to test the navigation ability of a large number of participants across many different countries. What we find is that there is no reliable connection between handedness and navigation ability. Then, we use our data to explore the prevalence of left-handedness across countries, and find that age, gender, and country of residency all have an effect on the ratio of left-handedness. Moreover, we find an effect of education on left-handedness in China, Indonesia, India, Taiwan, and Hong Kong.

## Introduction

The impact of handedness on cognition is a question of longstanding interest across several domains (Annett, 1998; de Kovel et al., 2019; Deep-Soboslay et al., 2010; Levy, 1976; Markou et al., 2017; Sartarelli, 2016). One of these domains concerns spatial cognition. A meta-analysis found that right-handers slightly outperformed others in spatial tasks such as mental rotation and picture assembly (Somers et al., 2015). However, the majority of the studies analysed tackled this question with few participants. In fact, the effect failed to reach significance when a single, large (sample = 210,916) study was excluded from the meta-analysis (Peters et al., 2006). This suggests that a robust association of hand preference with spatial cognition necessitates a large sample. Moreover, most studies have focused on small-scale spatial tasks (Cheng, Hegarty, and Chrastil 2020), rather than on large-scale spatial cognition. While right-handers have been found to outperform left-handers in small-scale spatial tasks (Krumina et al., 2018), the opposite effect was found in a large-scale task involving navigation (Piper et al., 2011).

The link between handedness and large-scale navigation is complicated by the fact that cultural differences have a significant impact on both. Differences in nationality and culture are associated with variation in spatial navigation ability (Coutrot et al., 2018; Spiers et al., 2021; Walkowiak et al., 2022; Newcombe et al., 2022). Hand-preference distribution also varies widely between countries, likely due to different cultural pressures (Papadatou-Pastou et al., 2020): left-handers constitute 15-20% of the population in North America but only 0.06-2.8% in China (Kushner, 2013; Raymond & Pontier, 2010; Zverev, 2006).

Both handedness and navigation ability vary not only across, but also within, populations. Previous demographic studies have shown a higher proportion of males use their left-hand, and studies of navigation ability suggest a male advantage in spatial tasks (Coutrot et al., 2018; Medland et al., 2017; Papadatou-Pastou et al., 2008). Part of this difference might relate to handedness. For example, a study of 879 people found left-handed males and right-handed females outperformed other-handed people of their respective genders in a mental rotation task (Sanders et al., 1982). However, studies using similar paradigms completed by other groups have shown the opposite effect (Yen, 1975; Gordon & Kravetz, 1991). Alternatively, it might be that the purported association between handedness and spatial ability is due to the association between gender and spatial ability, given that males are more likely to be left-handed. Hand preference also varies depending on age, perhaps reflecting a change in tolerance towards left-handers over time, as older people are more likely to have been forced to switch handedness (de Kovel et al., 2019; Kushner, 2013).

A further reason for clarifying whether handedness is associated with navigation performance concerns the development of diagnostic tools for dementia (Cogné et al., 2017). A recent study found that right-handed Alzheimer’s disease patients were almost three times less likely to have early onset disease than left-handed patients, but the connection between handedness and dementia is also far from clear (Ryan, Kreiner & Paolo 2020). Navigation ability has been shown to be an important predictor of dementia (Coughlan et al., 2018) and pre-clinical risk (Coughlan et al., 2019), and screening for handedness is a quick procedure that does not involve language translation, so if handedness is associated with navigation ability, the finding would support the inclusion of handedness in cross-cultural diagnostics. Finally, if there are differences in spatial ability associated with hand preference, this would have consequences for experimental design: left-handers are routinely excluded from brain imaging studies (Knecht et al., 2000; Vogel et al., 2003; Willems et al., 2014). However, if handedness is associated with spatial navigation ability, the exclusion of 10% of the population potentially leaves an aspect of spatial cognition underexplored.

The challenge in addressing the link between handedness and spatial ability is twofold. First, it is difficult to test for large-scale spatial ability (i.e. navigation) with a method that is predictive of real-world performance. Second, the impact of culture on both handedness and spatial ability, compounded with the potentially small effects, means that tackling the question would require a large sample size. Even when testing the link between handedness and small-scale spatial ability, the majority of existing studies have had relatively small sample sizes. This means factors such as age and gender might not be adequately accounted for, which potentially explains the divergence in results across studies (Lawton et al., 2016).

Here, we overcame past limitations by using Sea Hero Quest, an ecologically-valid navigation task (Coutrot et al. 2019), to test a diverse worldwide sample of individuals from 41 countries. The main aim of the study is to establish whether handedness has an impact on spatial ability. Additionally, we wanted to determine the distribution of hand preference across nations, clarify how it connects to sociodemographic factors such as age, gender and education, and explore if those sociodemographic factors mediate the relationship between handedness and spatial ability.

## Methods

### Participants

#### Demographic analysis

3,881,449 people played at least one level of Sea Hero Quest. Participants that had not entered all of their demographics were excluded from this study as were participants who were over 70 years old, a group with strong selection bias, which has previously been shown to result in increased performance (Coutrot et al., 2018). Only countries with at least 1000 players were included in our sample. This resulted in an analytic sample of 749,037 participants (390,732 males) from 58 countries with a mean age = 38.64 (*SD =* 14.56). 535,325 received tertiary education (university or college), 213,389 received secondary education or lower. 74,444 were left-handers (9.94%).

#### Spatial ability analysis

Starting from the same analytic sample as the demographic analysis, only participants who had completed at least eleven levels of the game were included in the analysis. This ensured a reliable estimate of spatial navigation ability in our analytic sample. This resulted in an analytic sample of 422,772 participants (226,087 males) from 41 countries with a mean age = 37.81 (*SD =* 14.17). 42,232 were left-handers (9.99%).

## Materials

Sea Hero Quest is a mobile video game that measures spatial navigation ability (Coutrot et al., 2018; Spiers, Coutrot & Hornberger, 2021). In wayfinding levels (45 levels out of a total of 75 levels), Sea Hero Quest asks participants to view a map featuring both their current position and their goal locations (Figure 1a). Participants are then asked to navigate a boat as quickly as possible towards goal locations in a specified order (Figure 1b). We selected a subset of 4 wayfinding levels of increasing but moderate difficulty appearing quite early in the game (levels 6, 7, 8, and 11), alongside two training levels (levels 1 and 2). Consent for the study was provided by the UCL ethics board, and informed consent was provided within the app.

**Fig. 1.**
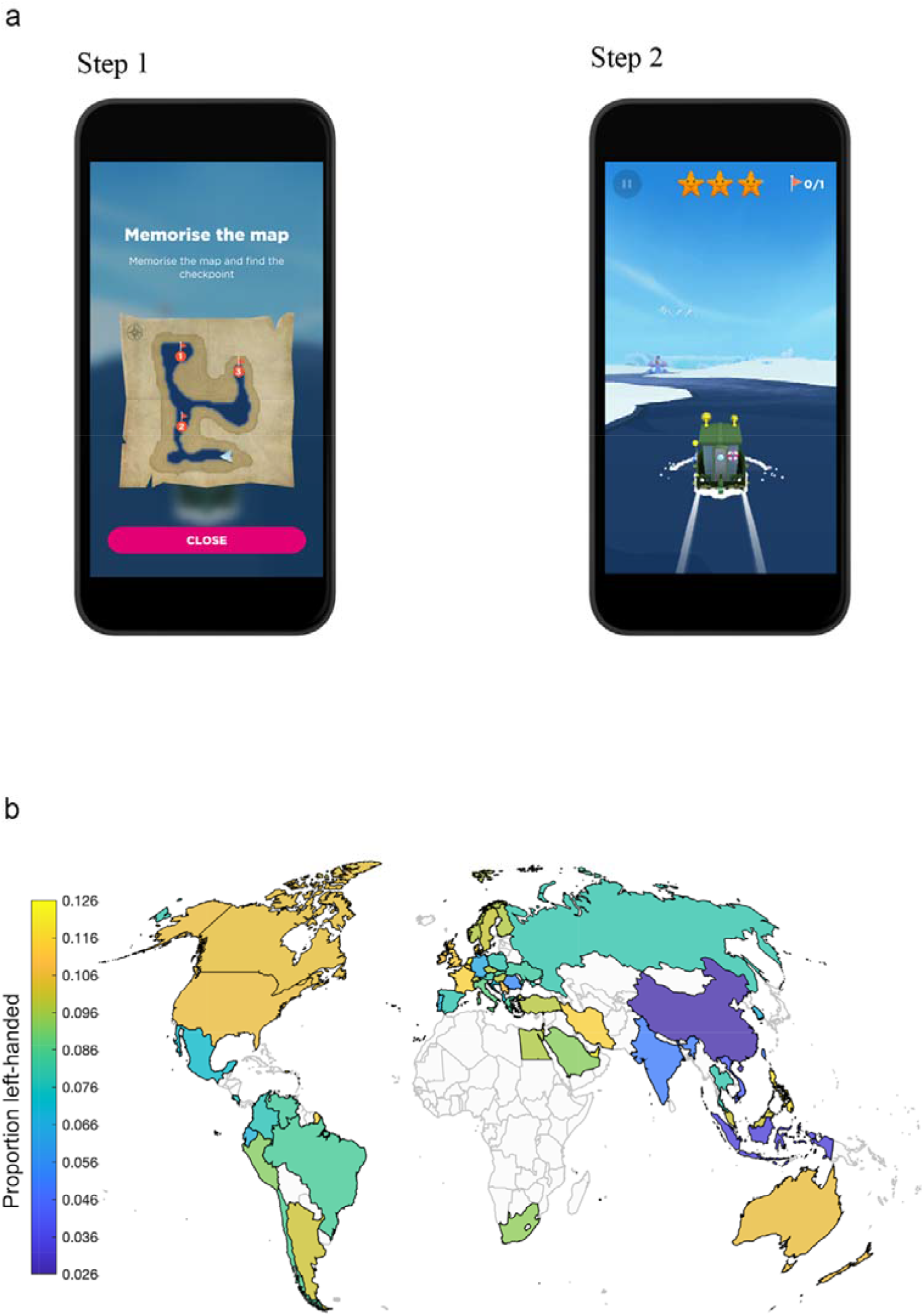
**(a)** Wayfinding task in the Sea Hero Quest app. Images show example screenshots from the game as they would appear on a mobile device. Step 1 involved viewing a map of the environment indicating the layout, current location (arrow) and checkpoints to navigate to in a given order. In the example above (level 11) the 3 checkpoints were used. Across game levels these varied from 1 to 5. After viewing the map, participants pressed the ‘close’ icon and the task transition to step 2. In step 2 participants taped left and right of the boat to steer it to the checkpoints and could swipe up to speed up or swipe down to slow the boat. **(b)** Map of left-handedness rate across countries.

Participant performance was measured by the Euclidean distance travelled in each level. The coordinates of their trajectories were sampled at Fs = 2Hz. We normalised the trajectory length in each level by the sum of the distances travelled over tutorial levels 1 and 2. These tutorial levels did not require any spatial ability and were designed to measure participants’ ability to control the virtual boat. We defined the wayfinding performance metric (WF_perf) as the first component of a principal component analysis over the normalised trajectory lengths of the four wayfinding levels included in the analysis (as in Coutrot et al., 2018).

Participants indicated handedness by selecting a hand on either the left or right side of the screen before they began the game. For analysis, age was from 19 to 69, gender had 2 classes (males, females), handedness had 2 classes (left, right) and education had 2 classes (up to secondary, tertiary).

## Results

### Demographics

We fit a multilevel logistic regression model with handedness as the response variable, age, gender, and education as fixed effects, and country as random effect (handedness ∼ age + gender + education + (1 | country)). All dependent variables had a significant effect on handedness: age (F(1,748708) = 464.45, p<0.001), gender (F(1,748708) = 925.56, p<0.001), and education (F(3,748708) = 48.435, p<0.001). The standard deviation of the country random effect was 0.34, 95% CI [0.28 0.41]. The variance partitioning coefficient (VPC), i.e. the proportion of observed variation in handedness that is attributable to the effect of clustering by country, is 3.61%.

The incidence of left-handedness in our sample was 9.94% and was smaller in females (8.95%) than in males (10.85%). It decreased with age (10.76% at 19 y.o. versus 8.68% at 70 y.o., Figure 2c) and with level of education (9.82% in participants with tertiary education, 10.25% with secondary education or lower, Figure 2b). Looking across countries, the gender effect is fairly consistent, with the exceptions of India, Indonesia, Costa Rica, and Saudi Arabia, where females are slightly more likely to use their left hand than males, Figure 2a. The Netherlands has the highest left-handers rate (12.95% left-handers), while China has the lowest (2.64% left-handers). The increase in left-handedness in participants with lower levels of education was mainly driven by China, Indonesia, India, Taiwan, and Hong Kong (N=26,223), where there is a lower rate of left-handers and where the education effect was much stronger than in the other included countries (N=722,814), see Figure 2b. In China, Indonesia, India, Taiwan, and Hong Kong, the average difference in left-hander ratio between participants with and without tertiary education was 4.49% (Chi2=96.74, p<0.001), while in the rest of the world it was 0.15% (Chi2=3.54, p=0.06).

**Fig. 2.**
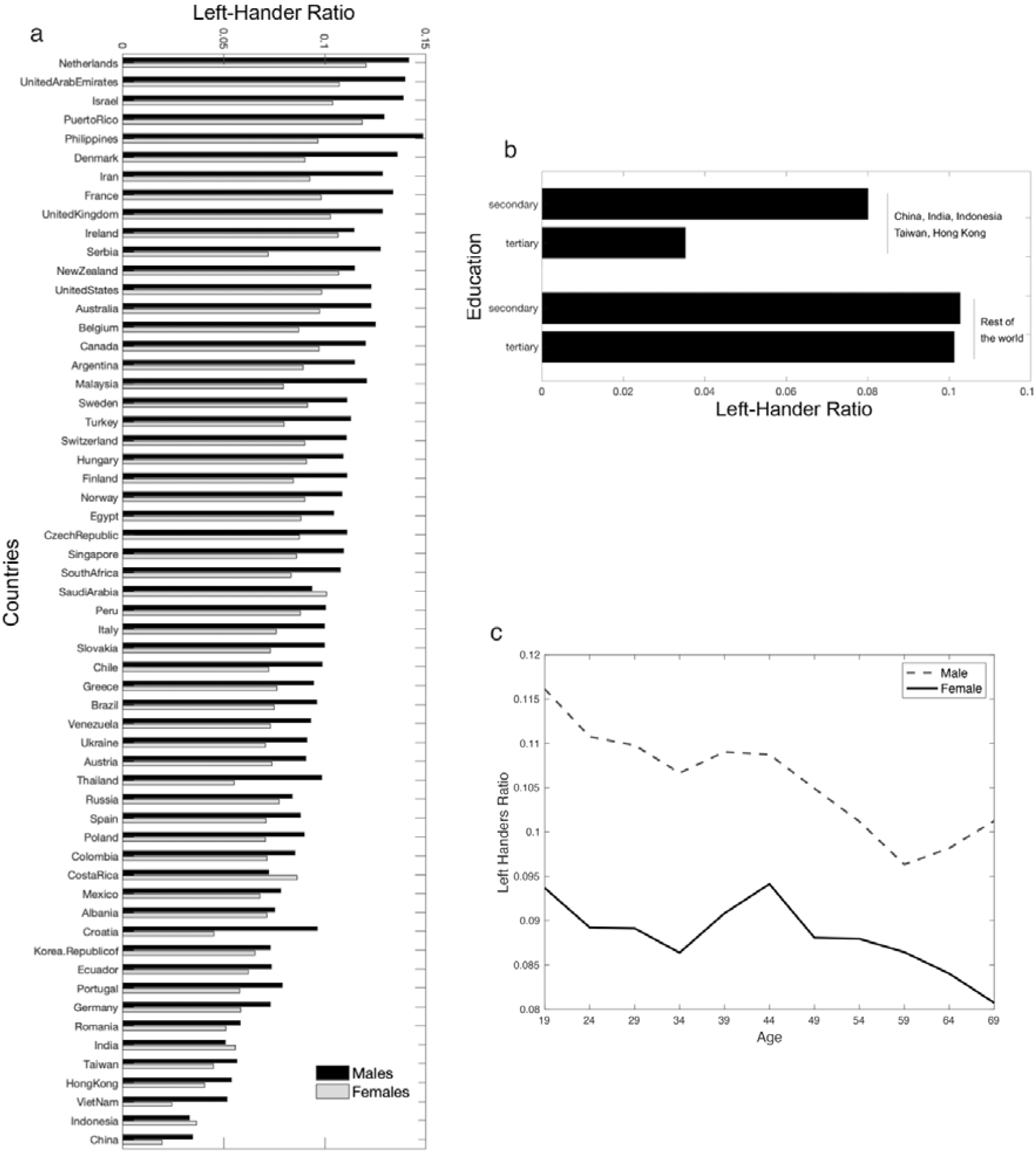
**(a)** Left-Handers ratio across countries, for males and females. **(b)** Left-Handers ratio in participants with tertiary education and with secondary education or lower. The first two bars correspond to participants from China, India, Indonesia, Taiwan, and Hong Kong, which have the lowest left-handers ratio. The second two bars correspond to participants from the other countries. **(c)** Left-Handers ratio across age and gender.

### Spatial ability

We fit a multilevel linear model with wayfinding performance as the response variable, handedness, age, gender, and education as fixed effects, with random slopes for handedness clustered by country (WF_perf ∼ age + gender + education + handedness + (handedness | country)). Age (F(1,422767) = 69094, p<0.001), gender (F(1,422767) = 23308, p<0.001), and education (F(3,422767) = 514.77, p<0.001) had a significant effect on wayfinding performance. In contrast, handedness did not have a significant effect (F(1,422767) = 1.72, p = 0.19). We measured the effect size of handedness on wayfinding performance with Hedge’s g. Overall, g = 0.045, 95% CI=[0.035, 0.055] (in females g=0.024, 95% CI=[0.008, 0.039], in males g = 0.027, 95% CI = [0.014, 0.04]), positive values corresponding to better performance in left-handers. As a point of comparison, for gender, g = 0.44, 95% CI = [0.43, 0.45], positive values corresponding to better performance in males. The standard deviation of the handedness effect across countries was 2.9e-3, 95% CI=[1.3e-4, 6.4e-2], which was very small compared to the residual standard deviation (0.80, 95% CI=[0.79, 0.80] and suggests that the differences of the handedness effect size between countries are negligible. This is illustrated in Figure 3a which shows the handedness slopes for each country. We see that there is very little variation across countries. To visually compare the magnitude of the effect of handedness to the effect of gender, the x-axis limits are set to the maximum values of the gender slopes across countries (−0.1,0.1).

**Fig. 3.**
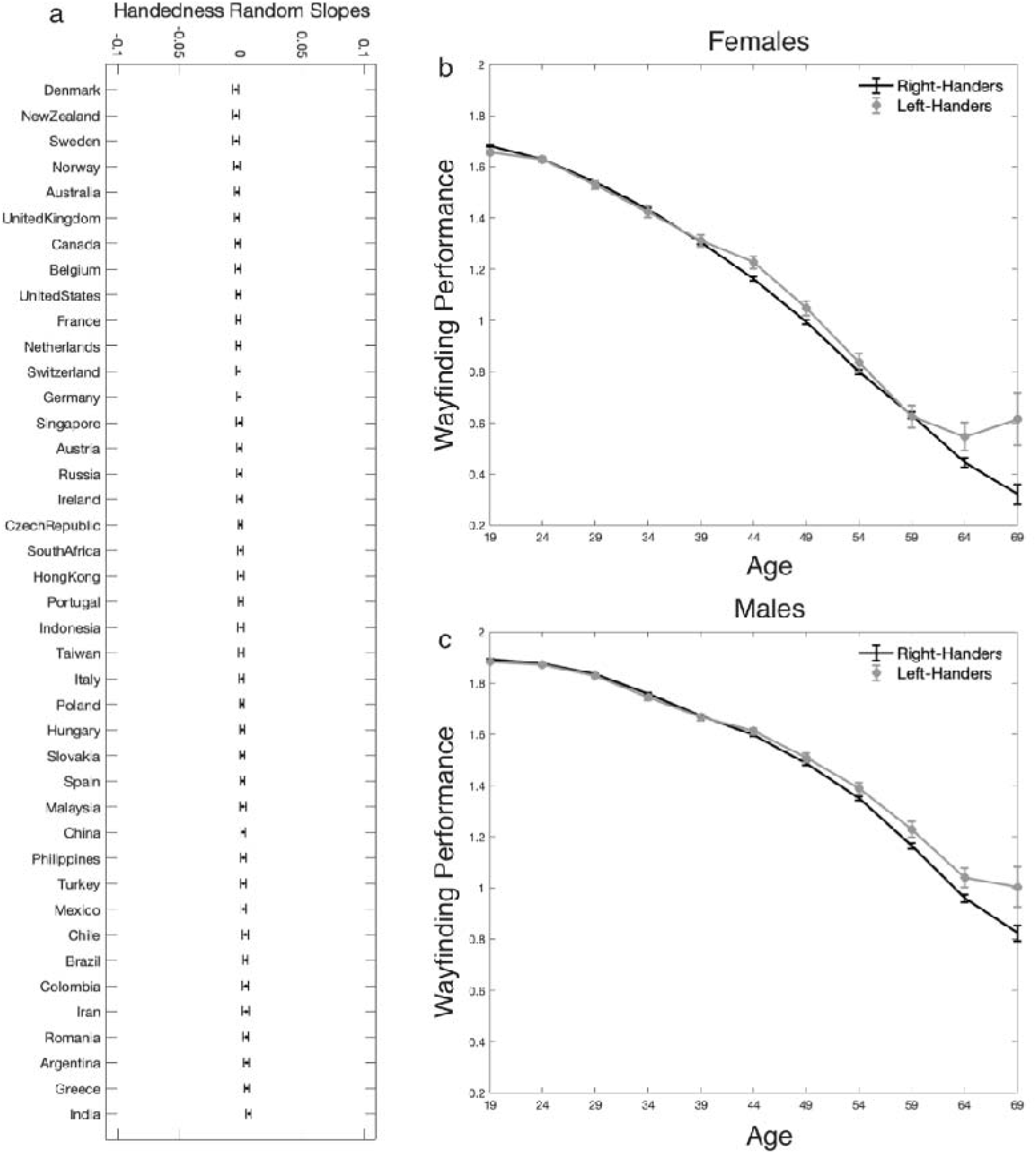
Navigation performance compared across countries, age, and gender. **(a)** Effect of handedness across countries. We fit a linear mixed model for wayfinding performance, with fixed effects for age, gender, education, and handedness, and random effect for country. We plot the handedness slopes for each country. To visually compare the magnitude of the variation of the handedness effect across countries to the variation of the effect of gender across countries, the x-axis limits are set to the maximum values of the gender slopes across countries. **(b-c)** Effect of handedness on wayfinding performance across age for males and females. The wayfinding performance has been averaged within 5-years time windows. We consider the apparent increase in performance in later life to be derived from a selection bias and plot data up to the age of 69, see Fig 4 for our analysis of the selection bias. Error bars correspond to 95% confidence intervals.

Figures 3b and 3c show the effect of handedness on wayfinding performance across age for males and females, respectively. Across the life course there is no difference in performance between left and right handers for males and females. A small but growing gap in performance appears for participants over 64 y.o., with left-handers outperforming right-handers (note that wayfinding performance has been averaged within 5-years time windows). This gap is most likely due to selection bias. Previous analyses using SHQ found a strong selection bias in older participants, which caused their wayfinding performance to be significantly higher than expected when compared to unselected participants of a similar age (Coutrot et al., 2018; Coutrot et a., 2022a,b). Figure 4 provides a further analysis of the issue, which supports the view that the divergence in performance between left and right handers in older age is due to selection bias.

**Fig. 4.**
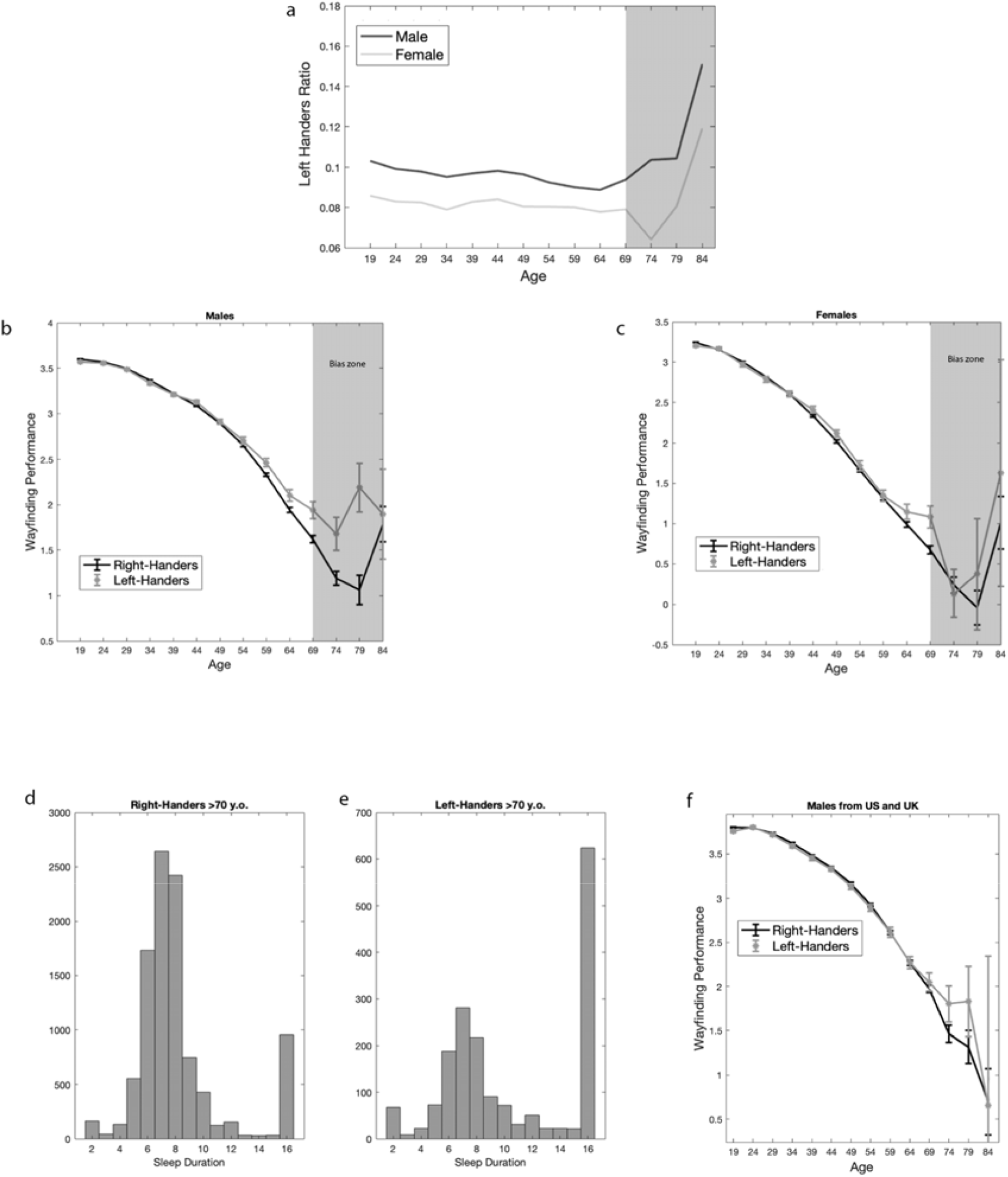
Analysis of selection bias. **(a)** Left-Handers ratio across age, for males and females. The ratio changes sharply after 70 y.o. The area in question is shadowed in the figure and marked as the “bias zone” **(b, c)** Effect of handedness on wayfinding performance across age for males and females. The wayfinding performance has been averaged within 5-years time windows. Error bars correspond to 95% confidence intervals. **(d, e)** Distribution of sleep duration for over 70 y.o. right-handers and left-handers. Those reporting to be left-handed show a substantive increase in reported sleep duration of over 16 hours a day for the group over 70, which is a pattern not predicted from lab studies (Coutrot et al., 2022b). **(f)** Effect of handedness on wayfinding performance across age for males from US and UK, a population we have found less selection bias in (Coutrot et al., 2018), which here similarly shows less effect of handedness on navigation with age than the global sample.

**Fig. 5.**
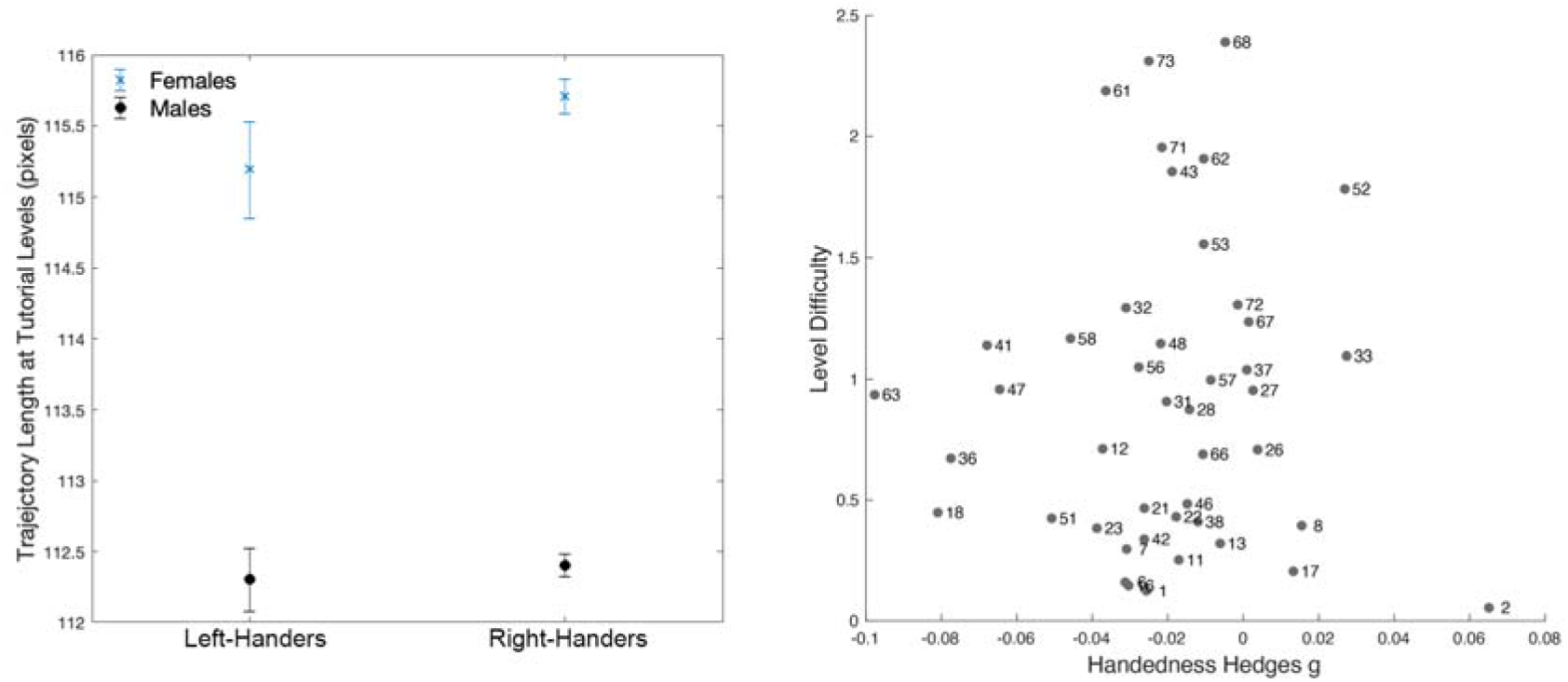
**(a)** Effect of handedness (Hedge’s g = -0.01) and gender (Hedge’s g = 0.11) on visuomotor skills. Visuomotor skills are estimated by the trajectory length at the tutorial levels (levels 1 and 2), which did not require any spatial ability. Error bars correspond to 95% confidence intervals. **(b)** Level difficulty as a function of Handedness effect size. Effect size has been estimated by Hedge’s g, with positive values corresponding to an advantage for the right-handers.

To verify whether handedness had an effect on visuomotor skills, we fit the same multilevel linear model as above, but with the trajectory length of the first two levels as the response variable. The first two levels were tutorial levels where large-scale navigation ability was not required, as the goal was visible from the starting point. Age (F(1,422767) = 5556.50, p<0.001), gender (F(1,422767) = 2323.90, p<0.001), and education (F(3,422767) = 286.15, p<0.001) had a significant effect on performance. On the other hand, handedness did not have a significant effect (F(1,422767) = 1.43, p = 0.23). Figure 4a shows the trajectory length at the first two levels for each gender and dominant hand. We measured the effect size of handedness on visuomotor skills with Hedge’s g. Overall, g = -0.014, 95% CI=[-0.024, - 0.004] (in females g=-0.019, 95% CI=[-0.034 -0.003], in males g = 0.002, 95% CI = [-0.011, 0.016]), positive values corresponding to better performance in left-handers. As a point of comparison, for gender, g = 0.11, 95% CI = [0.11, 0.12], positive values corresponding to better performance in males.

We tested whether the effect size of handedness was modulated by task difficulty. We selected a subset of participants who completed all Sea Hero Quest levels (75 levels, 10,626 participants) and computed Hedge’s g effect size between left-handers and right-handers in all wayfinding levels (N=44, not all Sea Hero Quest levels are wayfinding levels). We did not find a significant correlation between level difficulty and handedness effect size (Pearson’s correlation r = 0.02, p = 0.90, see Figure 4b). As in (Yesiltepe et al., 2022), we used the difference between the median trajectory length and the shortest trajectory length (better optimised) as a proxy for the level difficulty: Difficulty = (median(TL)-min(TL))/min(TL), with TL a vector containing the trajectory lengths of all participants at a given level.1

## Discussion

In this study, we examined the demographic data from 749,037 participants, across 58 countries and navigation performance from 422,772 from 41 countries. We found no reliable evidence supporting a benefit in spatial ability associated with hand preference, but positive evidence for an association with education and age on handedness prevalence. Here, we discuss handedness first in relation to spatial performance and then in terms of demographics.

Our findings challenge previous studies that suggested a significant relationship between an individual’s hand preference and their spatial performance in either small-scale (Krumina et al., 2018) or large-scale tasks (Piper et al., 2011). There are at least three reasons for this difference in findings: First, previous studies of spatial skill drew conclusions using small sample sizes. As a result, many studies were not designed to adequately address the research question, and those that were may have been susceptible to publication bias against null effects (Wilson & Bishop, 2018). Our study is the first to employ a large sample size to show a null effect of handedness on spatial ability, an approach that has been successful in other areas of research (e.g. null effects of bilingualism on executive tasks; Nichols et al., 2020). Second, in examining spatial navigation, we employed a mobile app with real-world ecological validity (Coutrot et al. 2019), whilst previous studies employed spatial visualisation tasks. Third, previous studies drew samples from single cultures, limiting the generalisability of their results. In the present study, we find our null effect to be universal across a broad span of cultures and languages.

Our use of large-population testing generates sufficient power to meaningfully explore the effects of potential moderating factors. Therefore, we examined whether an interaction between handedness and demographic properties impacted the effect of hand preference on wayfinding performance. We found that neither gender, nor age, nor the country, of our participants moderated the effect of handedness on spatial ability. In addition to large-scale navigation performance, we explored whether handedness might have impacted performance through visuomotor ability. However, as measured by our baseline test (distance in the tutorial), we find no evidence for this. We considered whether the effects of handedness only manifested in difficult tasks. But despite previous findings suggesting spatial granularity moderates the effect of handedness on spatial ability (as in Brunyé et al., 2012), we found the difficulty of our task did not have an effect either (for the effect of environmental difficulty on spatial tasks, see Sloane et al., 2015; He et al., 2021).

Demographically, we find an average of 9.94% left-handers overall, consistent with recent estimates (*10*.*6%*, Papadatou-Pastou et al., 2020). Like previous studies, we find more males report using their left hand compared to women (de Kovel, 2019; Peters et al., 2006). This gender difference is consistent across most countries, with only a few deviating from this pattern. We also find an overall decline in left-handedness with increasing age, as shown previously (Medland et al., 2004; McManus 2002). This finding may be due to a change in attitudes toward left-handedness (Kushner, 2013).

Additionally, our results show the ratio of hand preferences varies depending on the country. Only 2.6% of the participants from China were left-handed, a figure over 3 times smaller than the average for our sample. This finding is consistent with other studies showing that Chinese individuals are less likely than people from other countries to use their left hand. In this context, it has been suggested that attitudes toward left-handers are a proxy for tolerance towards difference more generally (Teng et al., 1976; Medland et al., 2004). While this finding may be partly due to attitudes towards conformity, results may also be influenced by the speed of industrialisation in China. In a country with a large influx of students who are the first in their generation to receive education, it may be more cost-effective to centralise resources and teach pupils to use the same hand in classrooms (Kushner, 2013). This is further evidenced by the effect of education. We found that in China, India, Indonesia, Taiwan, and Hong Kong, which have the fewest left-handers overall, people who had received tertiary education were less likely to be left-handed when compared to those who had received secondary education or less. In contrast, we found that education had no effect on the rest of the world (as found in Papadou-Pastou et al. 2020). This suggests that the fairly recent urbanisation of these countries may play a role in the incidence of left-handedness.

There are several limitations to this study. We use a self-reported measure, and in countries with negative attitudes towards left-handedness, participants might be reluctant to report being left handed. However, a meta-analysis found that self-reporting did not result in a statistical difference in left-handedness prevalence (Papadou-Pastou et al. 2020). Another limitation is the selection bias affecting older participants. Further work could examine this selection bias in more detail in an effort to elucidate why male left-handers have better spatial ability after 65 y.o. A hypothesis could have been that left-handers used to face increased educational difficulties. However, the absence of association between education and handedness in most countries (except China, India, Indonesia, Taiwan, and Hong Kong) does not support this hypothesis. Relatedly, by using a single-item measure of hand preference with icons to illustrate, we may not have captured the full spectrum of an individual’s handedness (Medland et al., 2004). Nevertheless, a longer questionnaire, such as the full Edinburgh Handedness Inventory, would not have been practical given our experimental paradigm (a mobile gaming application). Another limitation is that, while we draw from a truly international sample, we do not have representation from all countries. Nor can we ignore the cultural variation present within each country we sample and the lack of representation from more traditional societies (Medland et al., 2004). Future work along these lines would be valuable, given the relationship between the priority of particular skills (such as fishing over writing) and the cultural significance of handedness (Kushner 2013).

In conclusion, we provide a large sample of participants and countries to explore the impact of handedness on spatial ability. Our results demonstrate that across a large cross-cultural sample, hand preference is not associated with spatial ability. Moreover, our large sample allows us to verify that sociodemographic factors such as age, gender or education do not moderate the relationship between handedness and spatial ability. These results further our understanding of the interplay of handedness and cognition. They also have ramifications both in diagnostic screening and research design. Our study shows that left-handedness does not confer an increased general spatial ability, so in this respect, we found no support for including handedness in diagnostic screenings for dementia. And within the remit of navigation research, the null effect found in the present work allays the worries concerning the routine practice of excluding left-handers from brain imaging studies.

## Acknowledgements

Sea Hero Quest initiative funded and supported by Deutsche Telekom. Research was also supported by the Leverhulme Trust award DS-2017-026. Alzheimer’s Research UK (ARUK-DT2016-1) funded the analysis; Glitchers produced the game; and Saatchi and Saatchi London managed the creation of the game.

## Data Availability

A dataset with the pre-processed trajectory lengths and demographic information is available at https://osf.io/xfz8w/?view_only=08e221cfbff4416db02b1b2fda1b9539. The dataset with the full trajectories is available on a dedicated server: https://shqdata.z6.web.core.windows.net/. We also set up a portal where researchers can invite a targeted group of participants to play SHQ and generate data about their spatial navigation capabilities. Those invited to play the game will be sent a unique participant key, generated by the SHQ system according to the criteria and requirements of a specific project. https://seaheroquest.alzheimersresearchuk.org/ Access to the portal will be granted for non-commercial purposes.

## References

Annett M (1998): Handedness and cerebral dominance: The right shift theory. J Neuropsych 10: 459– 469

Brunyé, T. T., Gardony, A., Mahoney, C. R., & Taylor, H. A. (2012). Body-specific representations of spatial location. Cognition, 123(2), 229–239.

Cheng, Y., Hegarty, M., & Chrastil, E. R. (2020). Telling right from right: the influence of handedness in the mental rotation of hands. Cognitive Research: Principles and Implications, 5(1), 1–18.

Cogné, M., Taillade, M., N’Kaoua, B., Tarruella, A., Klinger, E., Larrue, F., … & Sorita, E. (2017). The contribution of virtual reality to the diagnosis of spatial navigation disorders and to the study of the role of navigational aids: A systematic literature review. Annals of physical and rehabilitation medicine, 60(3), 164–176.

Coughlan, G., Laczó, J., Hort, J., Minihane, A. M., & Hornberger, M. (2018). Spatial navigation deficits—overlooked cognitive marker for preclinical Alzheimer disease?. Nature Reviews Neurology, 14(8), 496–506

Coughlan, G., Coutrot, A., Khondoker, M., Minihane, A. M., Spiers, H., & Hornberger, M. (2019). Toward personalized cognitive diagnostics of at-genetic-risk Alzheimer’s disease. Proceedings of the National Academy of Sciences, 116(19), 9285–9292.

Coutrot, A., Silva, R., Manley, E., de Cothi, W., Sami, S., Bohbot, V. D., … & Spiers, H. J. (2018). Global determinants of navigation ability. Current Biology, 28(17), 2861–2866

Coutrot, A., Schmidt, S., Coutrot, L., Pittman, J., Hong, L., Wiener, J. M., … & Spiers, H. J. (2019). Virtual navigation tested on a mobile app is predictive of real-world wayfinding navigation performance. PloS one, 14(3), e0213272.

Coutrot, A., Lazar, A. S., Richards, M., Manley, E., Wiener, J. M., Dalton, R. C., … & Spiers, H. J. (2022). Reported sleep duration reveals segmentation of the adult life-course into three phases. Nature Communications, 13(1), 7697.

Coutrot, A., Manley, E., Goodroe, S., Gahnstrom, C., Filomena, G., Yesiltepe, D., … & Spiers, H. J. (2022). Entropy of city street networks linked to future spatial navigation ability. Nature, 604(7904), 104–110.

de Kovel, C. G., Carrión-Castillo, A., & Francks, C. (2019). A large-scale population study of early life factors influencing left-handedness. Scientific reports, 9(1), 1–11

Deep-Soboslay, A., Hyde, T. M., Callicott, J. P., Lener, M. S., Verchinski, B. A., Apud, J. A., … & Elvevåg, B. (2010). Handedness, heritability, neurocognition and brain asymmetry in schizophrenia. Brain, 133(10), 3113–3122

Gordon, H. W., & Kravetz, S. (1991). The influence of gender, handedness, and performance level on specialized cognitive functioning. Brain and Cognition, 15(1), 37–61.

He, Q., Han, A. T., Churaman, T. A., & Brown, T. I. (2021). The role of working memory capacity in spatial learning depends on spatial information integration difficulty in the environment. Journal of Experimental Psychology: General, 150(4), 666.

Knecht, S., Dräger, B., Deppe, M., Bobe, L., Lohmann, H., Flöel, A., … & Henningsen, H. (2000). Handedness and hemispheric language dominance in healthy humans. Brain, 123(12), 2512–2518.

Krumina, G., Skilters, J., Gulbe, A., & Lyakhovetskii, V. (2018, June). Effect of Handedness on Mental Rotation. In International Conference on Theory and Application of Diagrams (pp. 729–733). Springer, Cham.

Kushner, H. I. (2013). Why are there (almost) no left-handers in China?. Endeavour, 37(2), 71–81.

Lawton, C. A., Czarnolewski, M. Y., & Eliot, J. (2016). Laterality, sex, and everyday spatial behaviours: an exploratory analysis. Laterality: Asymmetries of Body, Brain and Cognition, 21(4-6), 745–766.

Levy, J. (1976). Cerebral lateralization and spatial ability. Behavior Genetics, 6(2), 171–188.

Markou, P., Ahtam, B., & Papadatou-Pastou, M. (2017). Elevated levels of atypical handedness in autism: Meta-analyses. Neuropsychology review, 27(3), 258–283.

McManus, C. (2002). Right hand, left hand: The origins of asymmetry in brains, bodies, atoms and cultures. Harvard University Press.

Medland, S. E., Perelle, I., De Monte, V., & Ehrman, L. (2004). Effects of culture, sex, and age on the distribution of handedness: An evaluation of the sensitivity of three measures of handedness. Laterality: Asymmetries of Body, Brain and Cognition, 9(3), 287–297.

Nichols, E. S., Wild, C. J., Stojanoski, B., Battista, M. E., & Owen, A. M. (2020). Bilingualism affords no general cognitive advantages: A population study of executive function in 11,000 people. Psychological Science, 31(5), 548–567.

Newcombe, N. S., Hegarty, M., & Uttal, D. (2023). Building a cognitive science of human variation: Individual differences in spatial navigation. Topics in Cognitive Science, 15(1), 6–14.

Papadatou-Pastou, M., Martin, M., Munafò, M. R., & Jones, G. V. (2008). Sex differences in left-handedness: A meta-analysis of 144 studies. Psychological Bulletin, 134(5), 677–699. https://doi.org/10.1037/a0012814

Papadatou-Pastou, M., Ntolka, E., Schmitz, J., Martin, M., Munafò, M. R., Ocklenburg, S., & Paracchini, S. (2020). Human handedness: A meta-analysis. Psychological Bulletin. Advance online publication. https://doi.org/10.1037/bul0000229

Peters, M., Reimers, S., & Manning, J. T. (2006). Hand preference for writing and associations with selected demographic and behavioral variables in 255,100 subjects: the BBC internet study. Brain and cognition, 62(2), 177–189

Piper, B. J., Acevedo, S. F., Edwards, K. R., Curtiss, A. B., McGinnis, G. J., & Raber, J. (2011). Age, sex, and handedness differentially contribute to neurospatial function on the Memory Island and Novel-Image Novel-Location tests. Physiology & behavior, 103(5), 513–522.

Raymond, M. & Pontier, D. Is there geographical variation in human handedness? Laterality 9, 35– 51, https://doi.org/10.1080/13576500244000274 (2004).

Ryan, J. J., Kreiner, D. S., & Paolo, A. M. (2020). Handedness of healthy elderly and patients with Alzheimer’s disease. International Journal of Neuroscience, 130(9), 875–883.

Sanders, B., Wilson, J. R., & Vandenberg, S. G. (1982). Handedness and spatial ability. Cortex, 18(1), 79–89

Sartarelli, M. (2016). Handedness, earnings, ability and personality. Evidence from the lab. PloS one, 11(10).

Slone, E., Burles, F., Robinson, K., Levy, R. M., & Iaria, G. (2015). Floor plan connectivity influences wayfinding performance in virtual environments. Environment and behavior, 47(9), 1024–1053.

Somers, M., Shields, L. S., Boks, M. P., Kahn, R. S., & Sommer, I. E. (2015). Cognitive benefits of right-handedness: a meta-analysis. Neuroscience & Biobehavioral Reviews, 51, 48–63.

Spiers, H. J., Coutrot, A., & Hornberger, M. (2021). Explaining World□Wide Variation in Navigation Ability from Millions of People: Citizen Science Project Sea Hero Quest. Topics in cognitive science.

Teng, E. L., Lee, P. H., Yang, K. S., & Chang, P. C. (1976). Handedness in a Chinese population: Biological, social, and pathological factors. Science, 193(4258), 1148–1150.

Vogel, J. J., Bowers, C. A., & Vogel, D. S. (2003). Cerebral lateralization of spatial abilities: a metaanalysis. Brain and cognition, 52(2),197–204.

Walkowiak, S., Coutrot, A., Hegarty, M., Velasco, P. F., Wiener, J. M., Dalton, R. C., … & Manley, E. (2022). Cultural determinants of the gap between self-estimated navigation ability and wayfinding performance: evidence from 46 countries. bioRxiv.

Willems, R. M., Van der Haegen, L., Fisher, S. E., & Francks, C. (2014). On the other hand: including left-handers in cognitive neuroscience and neurogenetics. Nature Reviews Neuroscience, 15(3), 193–201.

Wilson, A. C., & Bishop, D. V. (2018). Resounding failure to replicate links between developmental language disorder and cerebral lateralisation. PeerJ, 6, e4217.

Yen, W. M. (1975). Independence of hand preference and sex-linked genetic effects on spatial performance. Perceptual and Motor Skills, 41(1), 311–318.

Yesiltepe, D et al. (2022), Entropy and a Sub-Group of Geometric Measures of Paths Predict the Navigability of an Environment. Available at SSRN: https://ssrn.com/abstract=4170481

Zverev, Y. P. (2006). Cultural and environmental pressure against left-hand preference in urban and semi-urban Malawi. Brain and Cognition, 60(3), 295–303.

